# Weevil carbohydrate intake triggers endosymbiont proliferation: a trade-off between host benefit and endosymbiont burden

**DOI:** 10.1101/2022.07.06.498660

**Authors:** Elisa Dell’Aglio, Virginie Lacotte, Sergio Peignier, Isabelle Rahioui, Fadéla Benzaoui, Agnès Vallier, Pedro Da Silva, Emmanuel Desouhant, Abdelaziz Heddi, Rita Rebollo

## Abstract

Nutritional symbioses between insects and intracellular bacteria (endosymbionts) are a major force of adaptation, allowing animals to colonize nutrient-poor ecological niches. Many beetles feeding on tyrosine-poor substrates rely on a surplus of aromatic amino acids produced by bacterial endosymbionts. This surplus of aromatic amino acids is crucial for the biosynthesis of a thick exoskeleton, the cuticle, which is made of a matrix of chitin with proteins and pigments built from tyrosine-derived molecules, providing an important defensive barrier against biotic and abiotic stress. Other endosymbiont-related advantages for beetles include faster development and improved fecundity. The association between *Sitophilus oryzae* and *Sodalis pierantonius* endosymbiont represents a unique case study among beetles: endosymbionts undergo an exponential proliferation in young adults concomitant with the cuticle tanning, then they are fully eliminated. While endosymbiont clearance, as well as total endosymbiont titer, are host-controlled processes, the mechanism triggering endosymbiont exponential proliferation remains poorly understood. Here, we show that endosymbiont exponential proliferation relies on host carbohydrate intake, unlike the total endosymbiont titer or the endosymbiont clearance, which are under host genetic control. Remarkably, insect fecundity was preserved, and the cuticle tanning was achieved, even when endosymbiont exponential proliferation was experimentally blocked, except in the context of a severely unbalanced diet. Moreover, a high endosymbiont titer coupled with nutrient shortage, dramatically impacted host survival, revealing possible environment-dependent disadvantages for the host, likely due to the high energy cost of exponentially proliferating endosymbionts.

**Abstract Importance:** Beetles thriving on tyrosine-poor diet sources often develop mutualistic associations with endosymbionts able to synthesize aromatic amino acids. This surplus of aromatic amino acids is used to reinforce the insect’s protective cuticle. An exceptional feature of the *Sitophilus oryzae* / *Sodalis pierantonius* interaction is the exponential increase in endosymbiotic titer observed in young adult insects, in concomitance with cuticle biosynthesis. Here, we show that host carbohydrate intake triggers endosymbiont exponential proliferation, even in conditions that lead to the detriment of the host survival. In addition, when hosts thrive on a balanced diet, endosymbiont proliferation is dispensable for several host fitness traits. The endosymbiont exponential proliferation is therefore dependent on the nutritional status of the host, and its consequences on host cuticle biosynthesis and survival depend on food quality and availability.

## Introduction

Mutualistic symbioses are a powerful driving force for evolution, in that they promote adaptation to unfavorable environments^1^, allow the colonization of new ecological niches^2–8^, and prevent parasitic infections^9,10^. Insect colonization of poor nutritional substrates, such as cereals, plant sap, and blood, has been promoted by the establishment of trophic mutualistic symbioses with gut intracellular microorganisms, called endosymbionts^11^, that colonize the insect gut or form specific organs called bacteriocytes. Present in about 15% of known insect species^11^, the endosymbionts often provide an excess of nutrients to the host, thus allowing the insect to thrive even on unbalanced diet sources^12,13^, leading to an improvement of various fitness traits, such as fertility, lifespan, and stress tolerance^2,12,14^. In some cases, the endosymbionts have become essential for the host survival^14^.

The insect order of Coleoptera (the most diversified of all) is characterized by a thick exoskeleton, also called cuticle, constituted by proteins and pigments in a matrix of chitin that covers the whole body of the adult. The pigments that harden the cuticle use the tyrosine-derived 3,4-dihydroxyphenylalanine (DOPA) as a precursor^15^. Several beetles and weevils feeding on tyrosine-poor substrates, such as cereals, have acquired bacterial endosymbionts able to synthesize aromatic amino acids autotrophically and redirected a surplus of these amino acids to host cuticle biosynthesis^16–19^. Accordingly, aposymbiotic insects often show a paler cuticle as well as longer development and reduced fertility^5,17,18^. Other studies also stressed the ecological importance of the gut microbiota for cuticle reinforcement to enhance their protection from biotic and abiotic stress^9,16,20,21^.

On the endosymbiont side, one of the most common consequences of domestication is the loss of an independent lifestyle, often accompanied by a process of genome erosion^22^. This process is common in cereal-feeding weevils’ symbioses but is remarkable in the *Nardonella* endosymbiont of the red palm weevil *Rhynchophorus ferrugineus*, where the only metabolic pathway still present is the tyrosine synthesis pathway^17^.

In this context, the *Sitophilus oryzae* interaction with the endobacterium *Sodalis pierantonius* represents a unique case study. On one hand, it is the textbook model of mutualistic symbiosis with an obligate endosymbiont providing aromatic amino acids needed for building the host cuticle^23^, as well as other fitness advantages^5,13^. Notably, *S. pierantonius* has lost the ability to thrive outside bacteriocytes, and its titer among weevil’s strains are controlled by the host, as demonstrated by crosses between rapid-developing and slow-developing weevil strains, as well as by radiation experiments^24^. On the other hand, *S. pierantonius* has been recently acquired by substitution of a previous *Nardonella* symbiont (∼28K years), and the process of domestication is still ongoing. Indeed, *S. pierantonius* genome size is still comparable to the genomes of free-living bacteria such as *Escherichia coli*, and contains virulence genes such as components of a type 3 secretion system and the flagellum, although the process of pseudogenization is already quite advanced^25^. The expression of virulence genes is observed in particular during the insect metamorphosis when *S. pierantonius* endosymbionts leave the larval bacteriome where they were confined and migrate to the adult midgut, where they colonize new stem-cells at the apex of the insect caeca^26^ In terms of energy use, Rio and colleagues^27^ have shown the presence of a wide variety of *S. pierantonius* genes involved in polysaccharide catabolism. While the subsequent genome sequencing project has revealed that many of those are currently pseudogenized, the *malP* gene (involved in the degradation of maltodextrins), and a glucose-6-phosphate transporter are still predicted to be functional^25^. This suggests that the endosymbiont could potentially contribute to the catabolism of gut-assimilated carbohydrates.

Although wild endosymbiotic-free *S. oryzae* insects have never been found, they can be obtained in laboratory conditions by heat treatment^28^. This aposymbiotic lineage has a longer developmental time, reduced fecundity and a thinner, lighter cuticle with respect to the symbiotic animals^28,5^.

While in the majority of known symbioses between beetles and bacteria, a small population of gut endosymbionts is maintained throughout their whole lifespan, with little reduction in old beetles^17-19^, a striking feature of the *S. oryzae* / *S. pierantonius* symbiosis is that, right after metamorphosis, gut endosymbionts undergo an exponential proliferation phase, which is concomitant with the endosymbiont-dependent cuticle reinforcement. The gut endosymbiont exponential proliferation is followed by complete endosymbiont clearance driven by host apoptotic and autophagic mechanisms, allowing energy recycling from the bacteria to the host while avoiding inflammatory necrotic processes^23^. These mechanisms, which are triggered, at the transcriptional level, before the endosymbionts reach their higher titer, avoid tissue inflammation and the activation of the systemic immune response^29^. Since the endosymbiont exponential proliferation preceded the cuticle tanning process, it was suggested that such proliferation was necessary for the production of the excess of aromatic amino acids for cuticle biosynthesis, and hence beneficial for the host^23^. However, a causal link between endosymbiont exponential proliferation and cuticle tanning was not clearly established. Furthermore, a higher titer in aromatic amino acids was observed only in symbiotic weevils right after metamorphosis, suggesting that the differences in the nutritional status of symbiotic and aposymbiotic weevils also precedes the endosymbiont exponential increase.

Here, we asked whether the endosymbiont exponential proliferation observed in *S. oryzae* is controlled by the host, as the total bacterial titer^24^ and the clearance phase^23^, and if it is necessary for host cuticle tanning and fecundity. To answer this question, we have probed the plasticity of the endosymbiont dynamics by applying nutritional stress to the host and the endosymbiont, and by measuring the effects on endosymbiont dynamics, cuticle reinforcement, survival, and fecundity. Our results show that the endosymbiont exponential proliferation is triggered by carbohydrate provision, while its impacts on host fitness depend on food quality and availability.

## Results

### Endosymbiont exponential proliferation and endosymbiont-dependent cuticle tanning precede adult emergence

Females of *S. oryzae* lay eggs inside cereal grains, where the progeny develops up to early adulthood (**Fig. 1A**). After metamorphosis, gut endosymbionts are located in specialized cells, the bacteriocytes, at the apexes of gut caeca^26^, where they proliferate exponentially before a complete host-controlled clearance^23^. When adult weevils exit the grain by piercing a hole with their rostrum (*i*.*e*. emergence), the endosymbiont titer is close to its maximum, while the host cuticle tanning process, measured as a reduction in thorax redness^17,30^, is about to be completed (**Fig. S1**).

**Figure 1.**
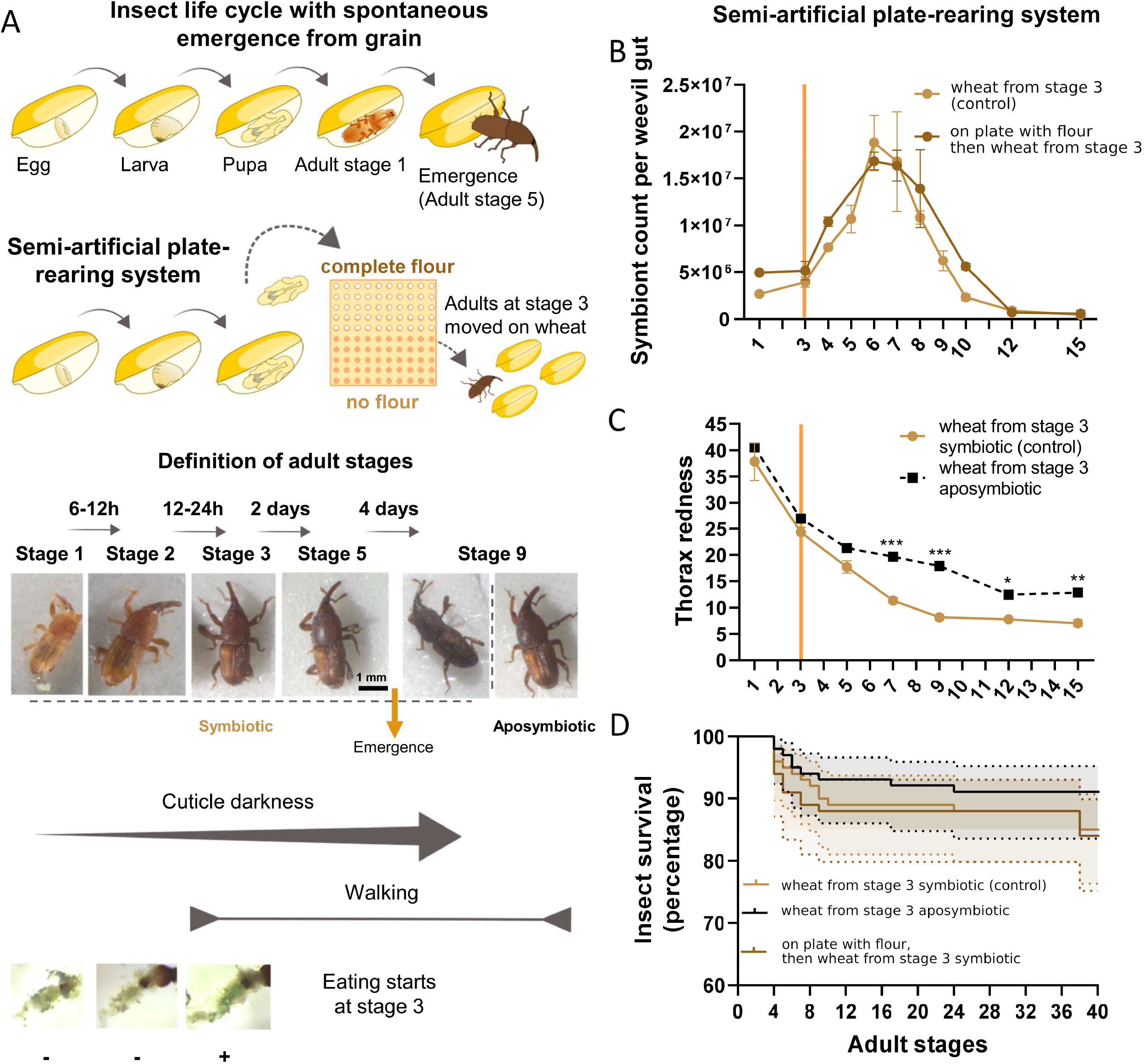
Endosymbiont exponential proliferation and endosymbiont-dependent cuticle tanning precede adult emergence. **A)** Schematic representation of the natural insect development on grain and the semi-artificial rearing system, after manual extraction of the pupae from the grains. Each pupa was kept on a well of a 10×10 well-plate (with whole wheat flour or without – control condition) until reaching adult stage 3, then kept on wheat grains. Insects were observed daily to monitor their development based on cuticle color, and ability to move and feed. Stage 9 aposymbiotic weevils are also shown for comparison. **B)** Endosymbiont dynamics of weevils reared on plates from the pupal stage with or without (control condition) whole flour supplementation until stage 3, then fed on wheat. **C)** Cuticle tanning progress, measured as a decrease in thorax redness, for plate-reared symbiotic and aposymbiotic weevils. **D)** Survival curves of aposymbiotic and symbiotic weevils reared as in **B)** and **C)**. Endosymbiont dynamics and cuticle comparisons were made by two-way ANOVA followed by Tukey’s multiple comparison test. Survival curves were analyzed with the Kaplan-Meier method followed by a Log-rank test. Shaded regions represent 95% CI. *: p < 0.05, **: p < 0.01; ***: p < 0.001. Error bars represent the standard error of the mean. Orange bars in **B)** and **C)** depict the day food was provided to control and aposymbiotic insects.

To tackle the mechanisms behind endosymbiont exponential proliferation, we have established a semi-artificial rearing protocol allowing timing and sampling of adults before and after grain emergence. Pupae manually extracted from grains were maintained on plates while daily monitoring their development (**Fig. 1A**). Adult stages were defined as follows: adults at stage 1 are orange-colored individuals unable to walk; adults at stage 2 (six to 12 hours after stage 1) are darker in color but still unable to walk; adults at stage 3 (12-24 hours after stage 2) are brown and mobile; all subsequent developmental stages are daily increments (**Fig. 1A**). From Stage 3 onwards, weevils were maintained in small groups of individuals (males and females) and fed with the appropriate dietary condition (the control condition is represented by whole wheat pellets). Flow cytometry quantification of endosymbiont titer showed that endosymbiont exponential proliferation started at stage 4, reaching its maximum at stage 6, while clearance was completed at stage 15 (**Fig. 1B**). In parallel, we observed the progressive darkening of the cuticle, which reached its maximum a few days after the higher endosymbiotic titer (stage 9, **Fig. 1C**), in agreement with weevils naturally reared on wheat grains (**Fig. S1**). From a comparison of both color and endosymbiont dynamics of plate-reared insects (**Fig. 1B-C**) and insects that naturally emerged from grains (**Fig. S1**), we identified stage 5.5 as the moment of emergence.

We confirmed that cuticle tanning was slower in aposymbiotic weevils (*i*.*e*. animals artificially depleted of endosymbionts by a heat treatment as in Nardon, 1973^28,5^), never reaching symbiotic levels (**Fig. S1B**: insects naturally emerged from grains; **Fig. 1C** plate-reared insects), even though adult survival was comparable between symbiotic and aposymbiotic insects in laboratory conditions (**Fig. 1D**).

In a similar way, insects which were laid on whole wheat flour pellets supplemented with a cocktail of antibiotics lacked the endosymbiont exponential proliferation phase (**Fig. S2A**) and showed an aposymbiotic-like cuticle (**Fig. S2B**). Although the endosymbiotic titer was reduced, a small endosymbiont population persisted in these insects, meaning that the antibiotic treatment was not successful in fully eliminating the endosymbionts. In agreement with that, the development length and the emergence rate of symbiotic insects fed on antibiotics were intermediate between those of control symbiotic and aposymbiotic weevils (**Fig. S2C and D**)^5,23^.

### Endosymbiont exponential proliferation is carbohydrate-dependent and detrimental to host survival when coupled with a nutrient shortage

Keeping pupae in whole wheat flour supplemented with E133 blue dye resulted in finding the blue dye in the gut of adult weevils from stage 3, suggesting that adult weevils start eating one day before endosymbiont exponential proliferation (**Fig. 1A**). In agreement with this finding, insects kept on plates without food from the pupal stage to adult stage 3, then moved to wheat grains, presented a similar endosymbiont dynamics as insects kept in whole wheat flour from the pupal stage to adult stage 3 (**Fig. 1B**), and no difference in insect survival was observed (**Fig. 1D**). In contrast, feeding adult weevils only from stage 4 or stage 5 onwards caused a delay in the endosymbiont exponential proliferation of two and three days, respectively (**Fig. 2A**), while the overall dynamic profile was unaltered. This suggests that nutrient provision is crucial to sustain endosymbiont exponential proliferation.

**Figure 2.**
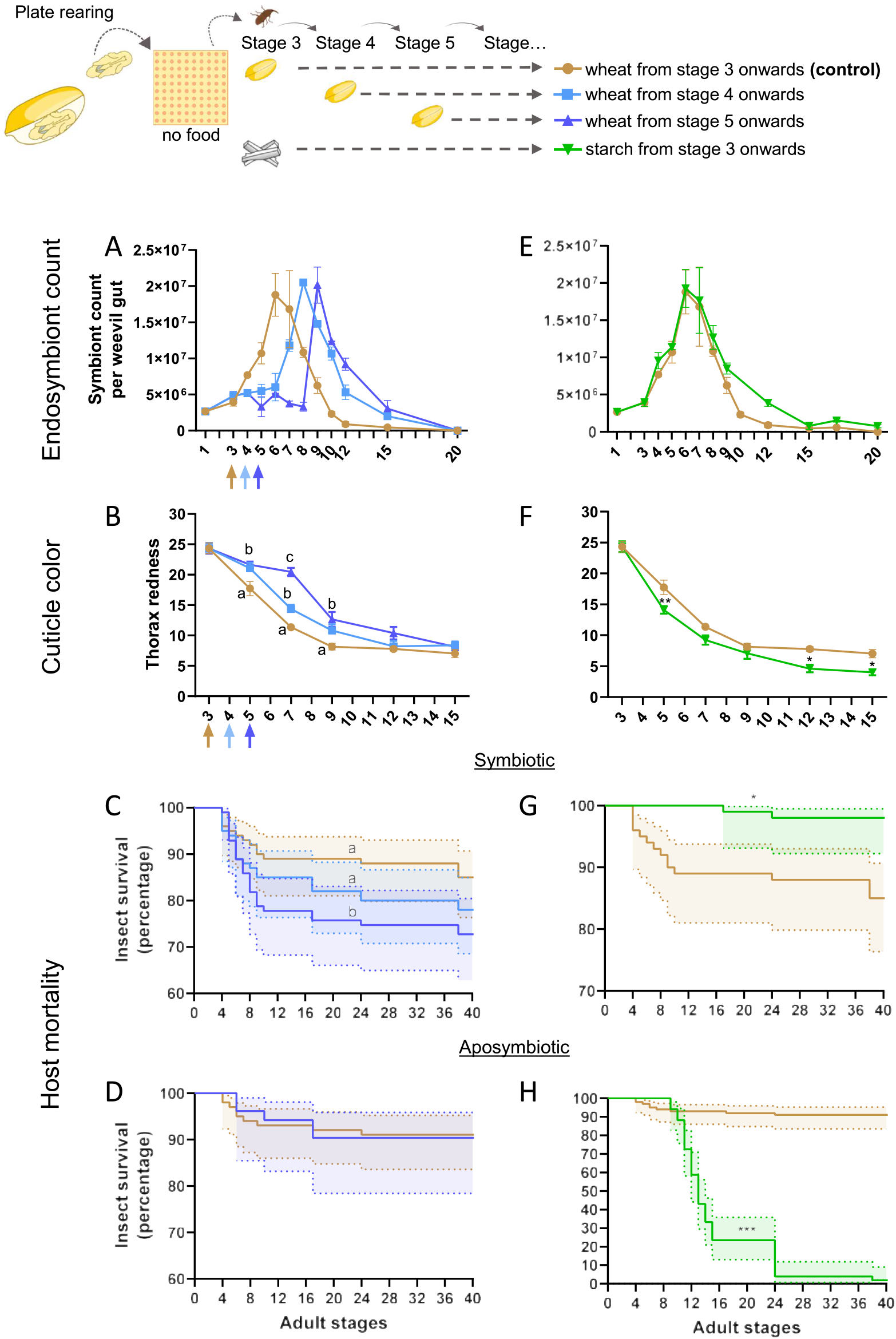
Endosymbiont exponential proliferation is carbohydrate-dependent and detrimental to host survival when coupled with nutrient shortage. Gut endosymbiont dynamics **(A)** and cuticle tanning progress **(B)** for symbiotic weevils fed with wheat grains from stage 3 onwards (control condition, as in **1B and 1C**, respectively), from stage 4, or stage 5. Colored arrows indicate the stage when food was administered. **C)** Symbiotic and **D)** aposymbiotic survival curves of weevils fed from stage 4 or 5 in comparison to control. Gut endosymbiont dynamics **(E)** and cuticle tanning progress **(F)** for weevils fed with starch grains from day 3 onwards, in comparison to control (as in **1B** and **1C**, respectively). **G)** Symbiotic and **H)** aposymbiotic survival curves of starch-fed weevils in comparison to control (as in **1D**). Endosymbiont dynamics and cuticle color comparisons were made by two-way ANOVA followed by Tukey’s multiple comparison test. Survival curves were analyzed with the Kaplan-Meier method followed by the Log-rank test. Shaded regions represent 95% CI. *: p < 0.05, **: p < 0.01; ***: p < 0.001. Error bars represent standard error of the mean. When more than one comparison is available, letters depict statistical significance between measures at each stage (panel B, thorax redness), or between survival curves (panel C).

We also observed a one-day delay in cuticle tanning of weevils fed from stage 4, and a three-day delay in weevils fed from stage 5 (**Fig. 2B**). Furthermore, the two-day delay in feeding caused a significant decrease in insect survival (**Fig. 2C)**, while no effect on fecundity (measured using the number of emerging descendants as a proxy) was observed (**Fig. S3A**). Interestingly, when starvation was applied to aposymbiotic weevils, no variation in survival rate was observed (**Fig. 2D**). However, when starvation was applied to symbiotic adults taken 15 days after emergence (when the gut endosymbiont population was already cleared, **Fig. S1A**), their survival was slightly higher than aposymbiotic animals (**Fig. S4**), thus suggesting either that aposymbiotic weevils are adapted to early adulthood starvation, or that the increased mortality in early symbiotic adults is due to the additional cost of maintaining the endosymbionts.

In a severely unbalanced diet, consisting only of starch (*i*.*e*. carbohydrates), endosymbiont dynamics was similar to the control condition (weevils fed with wheat from stage 3, **Fig. 2E**), and the cuticle tanning was completed faster (**Fig. 2F**), probably thanks to the higher friability of starch grains, which might be easier to break down and digest than wheat grains. A small endosymbiont gut population persisted until stage 27 (**Fig. S5**), suggesting that the host-controlled clearance^23^ can be delayed to extend endosymbiont presence in an extremely poor diet. Starch diet did not reduce but rather increased symbiotic insect survival in the first 40 days of adulthood (**Fig. 2G)**, although it completely abolished insect reproduction (**Fig. S3B**) and led to 100% mortality of aposymbiotic weevils (**Fig. 2H**), as previously described^23^.

DOPA accumulation was previously suggested as a putative molecular signal for endosymbiont clearance^23^. Here, the semi-artificial rearing system showed that DOPA increase was concomitant with the endosymbiont clearance rather than anticipating it (**Fig. S6A**), suggesting that DOPA is likely a transient molecule for mobilizing nitrogen-rich compounds freed by endosymbiont clearance. Indeed, DOPA increase was delayed by 2-3 days in weevils fed from stage 5 (**Fig. S6A**), coinciding with achieved cuticle tanning and the onset of endosymbiont clearance. In starch-fed weevils, DOPA levels resembled those of aposymbiotic weevils (**Fig. S6B**), suggesting that molecules enriched in aromatic amino acids are less abundant and/or recycled more efficiently than in grain-fed weevils to cope with the shortage of amino acids.

Overall, carbohydrate intake appeared necessary and sufficient to trigger endosymbiont proliferation. Higher mortality of young starved symbiotic insects (especially in comparison with aposymbiotic insects) points towards an excessive energy cost associated with the maintenance of the proliferating endosymbionts, or the development of a pathogenic behavior in the residual bacterial population (*e*.*g*. biofilm formation or bacterial migration outside of the caeca apexes). This suggests that, in contrast with the total endosymbiont titer^24^, and the subsequent clearance phase^23^, which are under host genetic control, the endosymbiont exponential proliferation in young adults depends on the host nutritional status.

### Endosymbiont exponential proliferation is dispensable for cuticle tanning and fecundity when coupled with a balanced diet

Since the majority of beetles relying on endosymbionts for cuticle tanning do not show an exponential increase in endosymbiont titer^16–19^, we asked whether a smaller endosymbiotic population would ensure cuticle tanning also in cereal weevils. To do so, we starved weevils at stage 4 and stage 5, after having fed them at stage 3. At first, endosymbiont exponential proliferation was triggered by feeding the weevils as control insects; next, the starvation phase arrested the endosymbiont proliferation, and triggered endosymbiont decrease from stages 5 to 8 (**Fig. 3A)**. Remarkably, a second exponential proliferation phase was observed after weevils were fed again, likely driven by carbohydrate intake, with a maximum titer at stage 11 and a complete clearance between stage 15 and 20 (**Fig. 3A**). In both exponential phases, the endosymbiont titer reached half the height of the control condition (weevils fed with wheat from stage 3), suggesting, as already demonstrated by genetic crosses^24^, that the total endosymbiont titer might be genetically controlled by the host. The first phase did not result in increased cuticle tanning, with stage 7 weevils still resembling control stage 4 weevils (**Fig. 3B**). In contrast, a few days after the second exponential proliferation phase, cuticle tanning was fully achieved (stage 12 **Fig. 3B**). Consistently with the hypothesis of a high metabolic cost associated with endosymbiont exponential proliferation, symbiotic weevils starved between stage 4 and stage 5 displayed 30% mortality, while no increase in mortality was observed for aposymbiotic weevils equally stressed (**Fig. 3C-D)**. Consistently with the hypothesis that DOPA accumulation represents a transient mobilization of nitrogen-rich storage molecules, two DOPA peaks were observed in concomitance with the two endosymbiont clearance phases (**Fig. S6C**).

**Figure 3.**
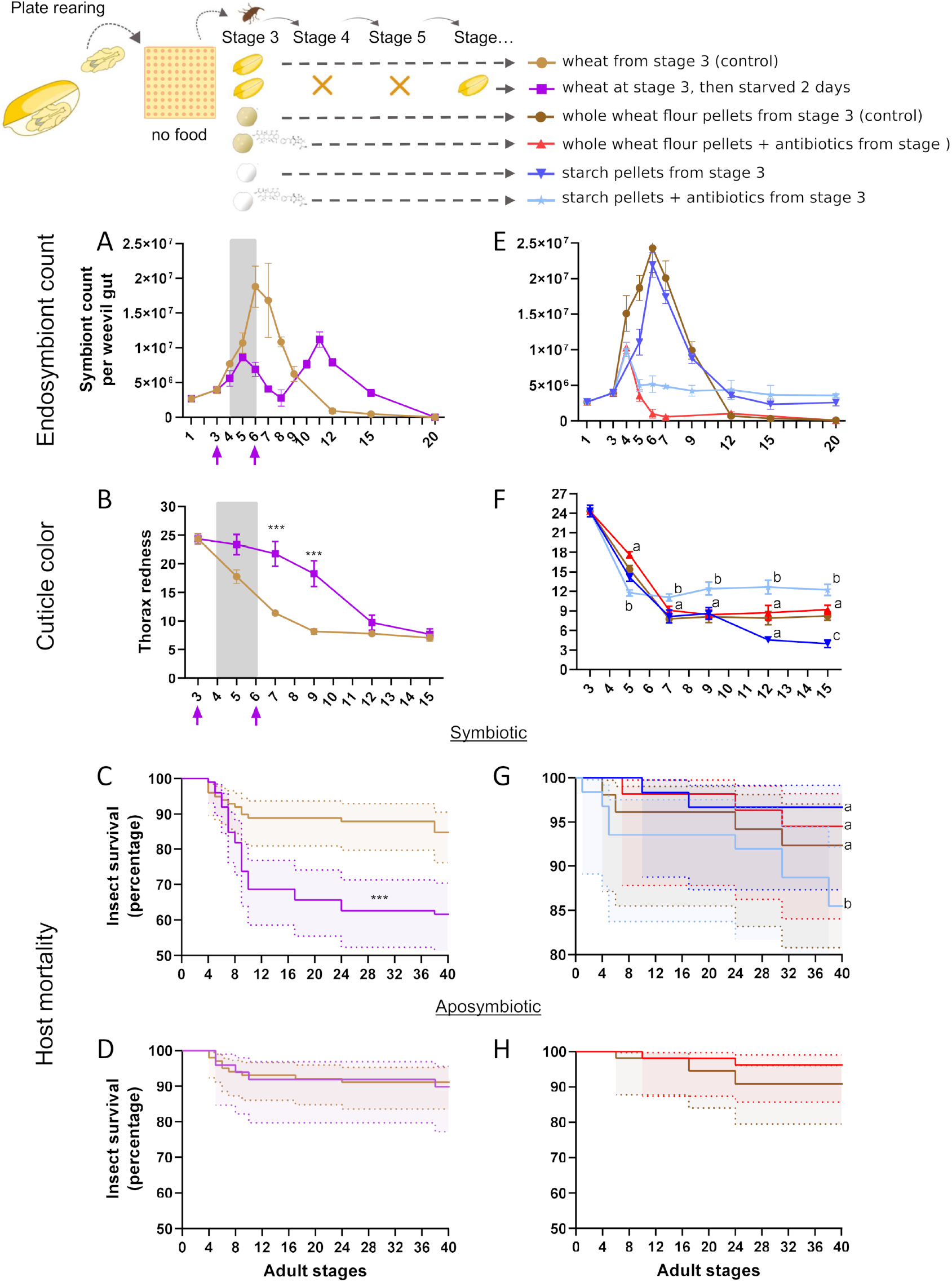
Endosymbiont exponential proliferation is dispensable for cuticle tanning and fecundity when coupled with a balanced diet. Gut endosymbiont dynamics **(A)** and cuticle tanning progress **(B)** for weevils fed with wheat at stage 3, then starved between stages 4 and 5, in comparison to control (as in **1B** and **1C**, respectively). Colored arrows indicate the stage when food was administered, and the gray area indicates the starvation period. **C)** Symbiotic and **D)** aposymbiotic survival curves of fed, then starved weevils in comparison to control (as in **1D**). Gut endosymbiont dynamics **(E)** and cuticle tanning progress **(F)** for weevils fed with wheat pellets supplemented or not (control condition) with antibiotics, or starch pellets supplemented or not with antibiotics from stage 3 onwards. **G)** Symbiotic and **H)** aposymbiotic survival curves of weevils fed with wheat pellets supplemented or not (control condition) with antibiotics, or starch pellets supplemented or not with antibiotics from stage 3 onwards. Endosymbiont dynamics and cuticle color comparisons were made by two-way ANOVA followed by Tukey’s multiple comparison test. Survival curves were analyzed with the Kaplan-Meier method followed by the Log-rank test. Shaded regions represent 95% CI. *: p < 0.05, **: p < 0.01; ***: p < 0.001. Error bars represent standard error of the mean. When more than one comparison is available, letters depict statistical significance between measures at each stage (panel F, thorax redness), or between survival curves (panel G).

We therefore hypothesized that a lower endosymbiont titer would be sufficient for cuticle tanning. To test this, while avoiding additional stress on the host, we fed adult weevils with whole wheat flour pellets supplemented with a cocktail of antibiotics^5,31^. While control weevils fed with whole wheat flour pellets from stage 3 displayed the expected endosymbiont exponential proliferation and clearance, the antibiotic supplementation triggered only a mild rise of endosymbionts at stage 4, followed by a complete clearance (**Fig. 3E**). With antibiotic supplementation, we did not observe differences in cuticle tanning (**Fig. 3F**), fecundity (**Fig. S3A**) or survival in symbiotic or aposymbiotic weevils (**Fig. 3G-H**), while DOPA accumulation was slightly reduced, likely due to the lower endosymbiont titer (**Fig. S6D**). The same endosymbiont dynamics was observed for weevils fed on starch pellets supplemented with antibiotics, except for the fact that, as already noted (**Fig 2E**), a small endosymbiotic population was retained longer (**Fig. 3E**). Remarkably, the antibiotic treatment combined with a severely unbalanced diet (starch only) reduced insect survival of starch-fed symbiotic animals (**Fig. 3G**), and their cuticle tanning was severely impaired: although faster (*i*.*e*. completed at stage 5) the process soon stopped at levels comparable to aposymbiotic weevils (**Fig. 3F**).

While these findings attested that endosymbiont exponential proliferation is highly energy-consuming and carbohydrate-dependent, they also revealed that, in the absence of a severely unbalanced diet, endosymbiont proliferation in young weevils is not required to ensure two of the major advantages known of this endosymbiosis, *i*.*e*. improved fecundity and cuticle tanning.

## Discussion

While previous studies have found that both the endosymbiont titer^24^ and the endosymbiont clearance in mature adults^23^ are controlled by the host, here we show that endosymbiont exponential proliferation in young adults relies on energy availability through the host diet, in the form of carbohydrates. Moreover, although previous results suggested that the exponential proliferation of *S. pierantonius* in *S. oryzae* young adults was necessary for ensuring enough building blocks for the adult cuticle biosynthesis, our new experimental approach has shown that, in the presence of a balanced diet constituted of whole wheat or whole wheat flour, the endosymbiont exponential proliferation phase is dispensable for both cuticle tanning and fecundity. In contrast, this increase in endosymbiont titer could be due to the host incapacity of controlling energy allocation to the endosymbionts, or to a transient virulent phase of the bacteria (*e*.*g*. biofilm formation, invasion of adjacent tissues and production of toxic metabolites), although we cannot exclude the presence of other fitness advantages for the host (*e*.*g*. stronger protection from parasites^10,20,32^) that could be detrimental in conditions of nutrient scarcity (**Fig. 4**).

**Figure 4:**
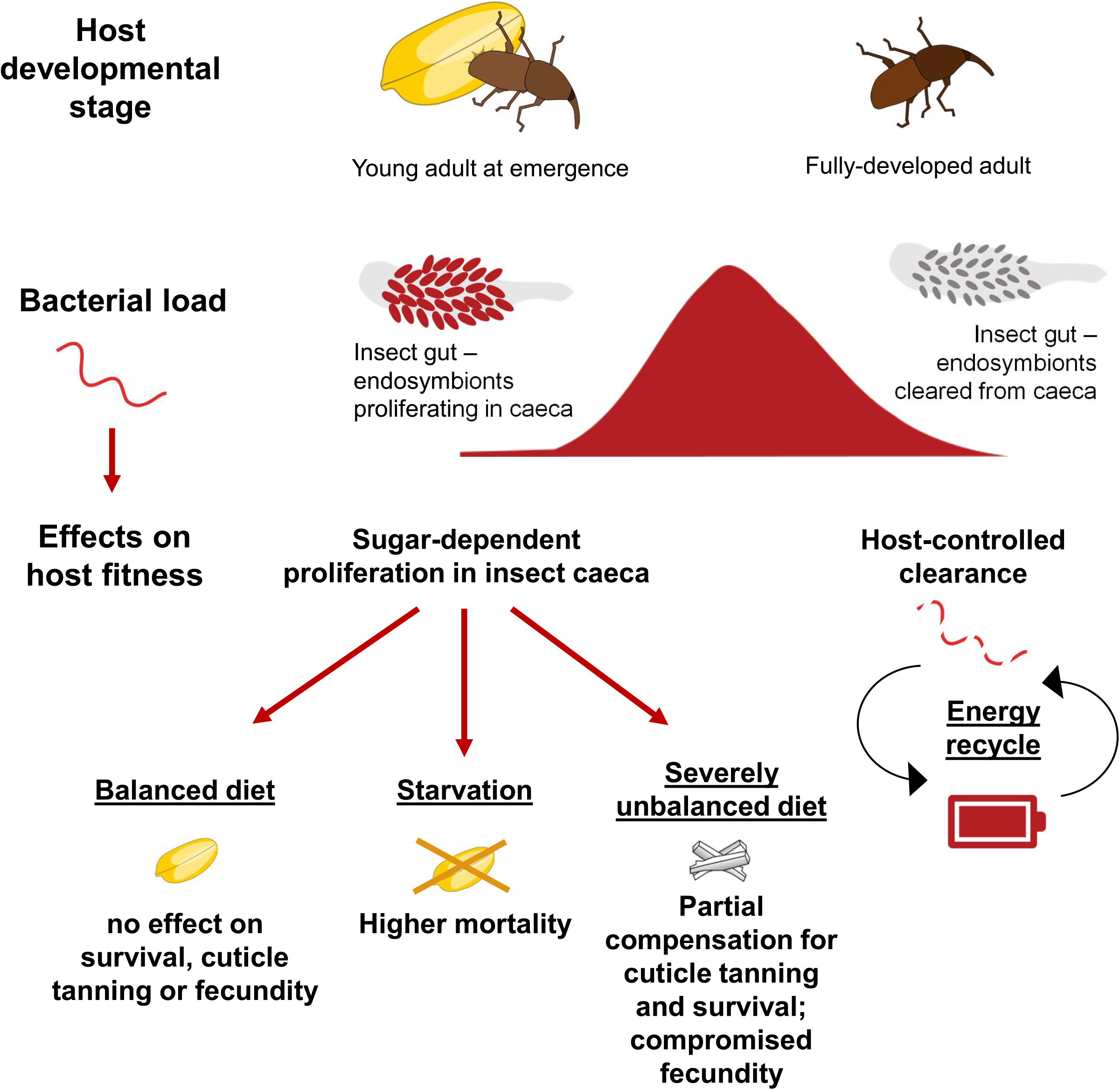
Schematic representation of various stages of the *S. oryzae/S. pierantonius* symbiosis and diet-dependent effects on host fitness traits. The outcome of endosymbiont exponential proliferation on host survival, cuticle biosynthesis and fecundity depends on nutrient quality and availability. Endosymbiont exponential proliferation is followed by host-controlled bacterial clearance that leads to energy recycling.

From an evolutionary perspective, growing evidence suggests the importance of the endosymbiont *S. pierantonius* for the acquisition of the modern *S. oryzae*’s life-style. The evolutionary origin of *S. oryzae* is still under debate, but growing evidence suggests that cereal-eating *Sitophilus* species originated from weevils inhabiting the cones of gymnosperms^33^, then, quite likely in concomitance with the development of agriculture^34^, the weevils transitioned to cereal stocks and co-evolved with domesticated cereal plants, likely helped by humans stocking cereals, acorns and nuts in close proximity. The presence of endosymbionts, and in particular the acquisition of *S. pierantonius* over the previous *Nardonella* ancestral endosymbiont within the Dryophoridae family^35-37^, seems to have been crucial for weevil adaptation to a substrate poor in proteins and vitamins. In general, agriculture has pushed towards bigger cereal kernels with higher starch content with respect to protein and vitamins^38-40^. In this context, the role of endosymbionts in complementing the host dietary needs seems to be even more relevant now than at the onset of agriculture, and might be beneficial in substrates poorer than wheat in proteins and vitamins, such as rice or white flour. The high abundance of starch in cereals, together with the fact that insects are still inside cereal grains at the onset of endosymbiont exponential proliferation, is likely the reason why the benefits of the endosymbiont exponential proliferation outweigh the associated costs.

At the same time, the starch-exclusive diet not only showed that a higher endosymbiont titer is advantageous in this case for cuticle tanning and survival, but also that a small endosymbiont population can be maintained after the clearance phase. Therefore, after the initial endosymbiont proliferation and clearance phases, *S. oryzae* weevils fed on starch resemble other more common beetle symbioses such as those of *O. surinamensis*^19^ and *R. ferrugineus*^17^. This supports the previous evidence^23^, by which the host plays a major role in orchestrating the endosymbiont clearance.

Symbiotic interactions are constantly evolving, in a continuum ranging from parasitism to mutualism, depending on changes in the interacting species and their environment^1,32, 41-43^. In insects, endosymbiont acquisition generally starts with domestication of parasites or commensals^43-45^. Host control over endosymbionts is often observed, as endosymbionts gradually lose the ability of autonomous life through genome shrinkage^46,47^ and point mutations^48^, sometimes retaining only the metabolic pathways that confer fitness advantage for the host^20^. Endosymbiont loss and/or replacement generally occur in concomitance with excessive genome shrinkage limiting host advantages^49–52^.

Other examples of host-controlled endosymbiont clearance during the host life cycle have been observed in obligate symbioses involving endosymbionts with extremely reduced genomes^53^. In contrast, hints of endosymbiont control over the host are rare and usually observed in facultative endosymbioses. For instance, recent findings have shown that free-living, facultative symbiotic algae are still able to colonize and proliferate in some species of cnidaria even when impaired in photosynthesis – the main host fitness advantage^54^. This raises the possibility of species-specific events of parasitic-like behavior of algal endosymbionts in contexts of nutrient shortage.

As in the algal-cnidaria case, this might be the consequence of a metabolic compatibility between the host and the endosymbiont and/or of the ability of the endosymbiont to efficiently use the allocated energy resources to proliferate^54,55^. Nutrient availability is another factor to consider, as endosymbionts thriving on limited energy resources are less likely to proliferate and trigger immune reactions. For instance, *Spiroplasma pulsoni* main energy source is constituted by lipid molecules which are scarce in the hemolymph of its host, *Drosophila melanogaster*, thus limiting *S. pulsoni* growth^56^. In contrast, the extremely high abundance of carbohydrates in the diet of adult *S. oryzae* likely favors endosymbiont proliferation. The precise timing of the event might also be linked to nutrient accessibility. Indeed, at larval stages, endosymbionts are confined in a larval bacteriome, and energy flux to this compartment is likely under strict host control, while the proximity of endosymbionts and the gut epithelium after endosymbiont colonization of adult gut caeca during metamorphosis might favor direct energy flux to the bacteria^26^.

Interestingly, *S. oryzae*’s sister species, *Sitophilus zeamais* presents similar endosymbiont dynamics, while the related species *Sitophilus granarius* shows lower and more constant endosymbiont levels in young adults^23^, suggesting different trajectories in coevolution for the control of endosymbiont titer. Comparison of different *Sitophilus* species and their endosymbionts in various stress conditions, together with artificial endosymbiont replacement and genetic modification strategies, would provide an ideal model for probing the mechanisms and constraints of endosymbiont domestication^57,58^.

## Material and Methods

### Insect rearing and growth conditions

The symbiotic *S. oryzae* population is constituted by a wild-derived strain (Azergues valley, Rhône, France), introduced into the laboratory in 1984 and maintained ever since. This strain contains exclusively the *S. pierantonius* endosymbiont. The aposymbiotic strain was obtained by heat treatment in 2010, following the protocol described in Nardon (1973)^28^. Aposymbiotic weevils were maintained alongside the symbiotic population ever since, in the same standard rearing conditions, in plastic boxes at 27 °C and 70% relative humidity, in the dark. Both strains (symbiotic and aposymbiotic) were routinely fed with organic wheat grains sterilized at −80 °C. Insects were kept in plastic boxes at 27 °C and 70% relative humidity, in the dark.

For antibiotic supplementation experiments, wheat flour pellets were prepared using commercial whole wheat flour (Francine, France) or starch (Stijfsel Remy, Belgium), with the addition of 0.1% (v/v) of chlortetracycline (Sigma-Aldrich) and 0.5% (v/v) penicillin G (Sigma-Aldrich). To prepare the pellets, flour/starch and, when needed, antibiotics were mixed with water (q.s.) to make a smooth dough. The dough was spread on a plastic surface and dried overnight at room temperature, then cut into little round pieces (pellets of ca. 5 mm in diameter) and stored at 4°C before use.

For analysis of insect development on antibiotic-supplemented with whole wheat flour pellets, two-week-old symbiotic and aposymbiotic adult weevils (n= 50) were fed for 24 hours with 20 whole wheat flour pellets (supplemented or not with antibiotics), then insects were removed and the pellets were kept in the incubator and observed daily to monitor: the day of progeny emergence, the number of emergents, the endosymbiont titer at emergence as well as the thorax cuticle color 12 days after emergence.

To monitor the moment adult weevils start eating after metamorphosis, pupae were manually extracted from grains and kept in plate wells with whole flour supplemented with E133 dye (100 μl of dye for 3.5 g flour). The E133 dye was first mixed with the flour with water (q.s.), let dry at room temperature overnight, and then ground. Guts of insects corresponding to various developmental stages (from stage 1 to stage 9) were dissected and observed with light microscopy.

### Endosymbiont quantification by flow cytometry

The protocol for endosymbiont quantification was modified from Login et al., 2011^59^. Briefly, a minimum of three pools (per condition and developmental stage) of four midguts each were dissected in TA buffer (25 mM KCl, 10 mM MgCl_2_, 250 mM sucrose, and 35 mM Tris/HCl, pH 7.5). The samples were manually ground in 100 μl TA buffer up to homogenization and centrifuged at 0.5 rpm for 2 minutes to sediment impurities. The supernatant was diluted in 400 μl TA buffer, then filtered with a 40 μm Flowmi filter (SP Scienceware) and centrifuged at 10 000 rpm for five minutes. The supernatant was discarded and the pellet was kept at 4°C in 4% paraformaldehyde (PFA, Electron Microscopy Science) before analysis.

Before quantification, pellets were centrifuged at 11 000 rpm for 20 minutes at 4°C, the PFA supernatant was discarded and samples were resuspended in 700 µl ultrapure water and 0.08% of SYTO9 dye (Invitrogen). additional water dilutions were made if the bacterial concentration was above the detection limit of the instrument.

Quantification was performed with BD Accuri C6 Plus cytometer (flow: 14 μl/min for 1 minute, cut off at 6000). Normalization was obtained by subtracting the values obtained from guts of aposymbiotic weevils. All measurements performed for endosymbiont titer analyses are independent from measurements of cuticle color, insect survival, fecundity, and DOPA quantifications.

### Analysis of cuticle color

The cuticle darkening process was monitored at various insect stages and conditions by using the Natsumushi software v. 1.10^17,30^ on pictures taken with an Olympus XC50 camera attached to a Leica MZFLIII binocular and the CellF software (Olympus Soft Imaging System) under the same lighting conditions. Quantification was performed as illustrated in Anbutsu et al., 2017^17^ using the thorax region, because of its color uniformity. Briefly, pixels with brightness over the top 10% or below the bottom 10% were excluded from the analysis. Then RGB values for all (= n) pixels were measured and averaged by Σ (R – mean [R, G, B])/n to obtain the proxy redness thorax mean value. Eight to 15 individuals were measured per condition and stage. All measurements performed for cuticle color analyses are independent from measurements of endosymbiont titer, insect survival, fecundity, and DOPA quantifications.

### Survival measurements

For plate-reared weevils, insects were isolated at the pupal stage on plate wells and assigned to a specific diet (n = 100 insects per diet condition, of mixed sexes, unsexed individuals) starting at adult stage 3, except for whole flour-reared weevils of Fig. 1B and Fig. 1D, which were reared on whole wheat flour from the pupal stage up to adult stage 3. Dead weevils were counted daily between stage 4 and stage 10, then weekly up to stage 40.

For weevils naturally emerging from grains, insects were isolated at emergence, kept for two weeks, and then starved for two days. Dead weevils were counted daily from the 16^th^ to the 22^nd^ day after emergence, then once per week up to the 44^th^ day after emergence.

All measurements performed for cuticle insect survival are independent from measurements of endosymbiont titer, color analyses, fecundity, and DOPA quantifications.

### Fecundity

We used the number of emerging descendants as a proxy to measure insect fecundity of plate-reared symbiotic and aposymbiotic weevils, as well as weevils fed from stage 5, starch-fed weevils, and antibiotic-fed weevils. Fifteen couples of randomly-paired male/female weevils per condition were established at stage 3. Then, weevils were subjected to the specific diet condition up to stage 8. At this point, antibiotic-fed weevils were shifted to the control diet (wheat grains). From stage 8 up to stage 45, the diet was changed every 3, 4, or 5 days (20 pellets each time). All wheat or starch grains were kept for two months to allow the emergence and counting of the progeny.

All measurements performed for fecundity analyses are independent from measurements of endosymbiont titer, cuticle color, insect survival, and DOPA quantifications.

### DOPA measurements

Measurements of free DOPA were performed on pools of frozen weevils (each pool made of three weevils) and the analysis was performed on three to five replicates per condition and stage. The whole weevil body was used for the analysis. Measurements were performed as in Vigneron et al., 2014^23^, using norvaline as an internal standard and a reverse phase HPLC method with a C18 column (Zorbax Eclipse-AAA 3.5 um, 150 × 4.6 mm, Agilent Technologies).

All measurements performed for DOPA analyses are independent from measurements of endosymbiont titer, cuticle color, insect survival, and fecundity.

## Acknowledgments

This work was funded by the ANR UNLEASh (ANR UNLEASH-CE20-0015-01 - R. Rebollo). We thank Aurélien Vigneron, Martin Kaltenpoth and Tobias Engl for interesting discussions.

## Authors contribution

Conceptualization: EDA, AH, RR; Methodology: EDA, VL, SP, IR, FB, AV; Writing: EDA, ED, AH, RR; Visualization: EDA, RR; Supervision: PDS, ED, AH, RR; Funding acquisition: RR.

## Supplemental information

**Figure S1. Endosymbiont dynamics and cuticle tanning in grain-reared weevils**. Gut endosymbiont dynamics **(A)** and cuticle tanning progress **(B)** for weevils developed in wheat grains and emerged spontaneously as adults. Results confirm and extend previous findings^23^. Red bar: emergence (E) from grain. All subsequent time points (E+X) represent days after emergence. Cuticle color comparisons between symbiotic and aposymbiotic weevils were made by two-way ANOVA followed by Tukey’s multiple comparison test. *: p < 0.05, **: p < 0.01; ***: p < 0.001. Error bars represent standard error of the mean.

**Figure S2: Symbiotic weevils on whole wheat flour pellets supplemented with antibiotics resemble aposymbiotic weevils**. A group of 100 symbiotic or aposymbiotic weevils was left for 24 hours with 30 whole wheat flour pellets supplemented or not with antibiotics. The weevils were then removed and the pellets kept for monitoring: **A)** endosymbiont titer at emergence, **B)** thorax redness at 12 days after emergence, **C)** day of progeny emergence from grain, and **D)** egg laying rate for symbiotic insects laid in whole wheat flour pellets (See STAR Methods) supplemented (red) or not (mustard) with antibiotics, compared to aposymbiotic insects fed in the same way. Comparisons were performed by Kruskal-Wallis test or t-test for experiments with only two populations. *: p < 0.05, **: p < 0.01; ***: p < 0.001. Error bars represent standard error of the mean.

**Figure S3: Comparison of insect fecundity under various stress treatments. A)** Couples of one male-one female eight-day-old weevils fed with wheat grains from stage 3 onwards (control condition), fed from stage 5, antibiotic-treated, or aposymbiotic weevils were reared on twenty wheat grains. The batch of grains was changed every 3 to 5 days until weevils reached 45 days of age. The effect of each condition on fecundity was calculated with a mixed-effect model with Geisser-Greenhouse correction followed by Tukey’s multiple comparison test (with fixed batch variable). *: p < 0.05, **: p < 0.01; ***: p < 0.001. Error bars represent standard error of the mean. **B)** Ovaries of stage 24 control weevils or weevils fed on starch from stage 3 onwards. The starch-fed weevils did not correctly develop the ovaries, and no eggs/progeny were retrieved from the starch pellets.

**Figure S4: Effect of starvation on adult weevils after endosymbiont clearance**. Symbiotic and aposymbiotic weevils at 15 days after emergence were either **(A)** kept on wheat grains or **(B)** starved for two days (E+16-E+17) before being fed again with wheat grains. The grey region represents the starvation period. Survival comparison between symbiotic and aposymbiotic weevils until E+44 was performed with the Kaplan-Meier method followed by the Log-rank test. Shaded regions represent 95% CI. *: p < 0.05, **: p < 0.01; ***: p < 0.001. Error bars represent standard error of the mean.

**Figure S5: Extended endosymbiont survival in the condition of a severe unbalanced diet**. Prolonged endosymbiont dynamic profile of starch-fed weevils from stage 3 onwards compared to control weevils (fed with wheat grains from stage 3 onwards). Note the retention of a small endosymbiont population until stage 25.

**Figure S6: In symbiotic weevils, free DOPA levels increase during endosymbiont clearance except in conditions of a severely unbalanced diet**. Free DOPA levels in the whole insect body were measured for weevils subjected to various stress treatments. **A)** control weevils (fed with wheat grains from stage 3) and weevils fed from stage 5; **B)** control weevils (as in **A**), aposymbiotic weevils, and weevils fed on starch from stage 3; **C)** fed, then starved weevils and control weevils (as in **A**); **D)** weevils fed on wheat or starch pellets supplemented or not with antibiotics. Here, the wheat pellet diet from stage 3 represents the control condition. Comparisons were performed by two-way ANOVA followed by Tukey’s multiple comparison test. *: p < 0.05, **: p < 0.01; ***: p < 0.001. Error bars represent standard error of the mean. When more than one comparison is available, letters depict statistical significance between measures at each stage (panel B and D).

