## Supplementary figures and images for "Weevil carbohydrate intake triggers endosymbiont proliferation: a trade-off between host benefit and endosymbiont burden"

### Figure S1

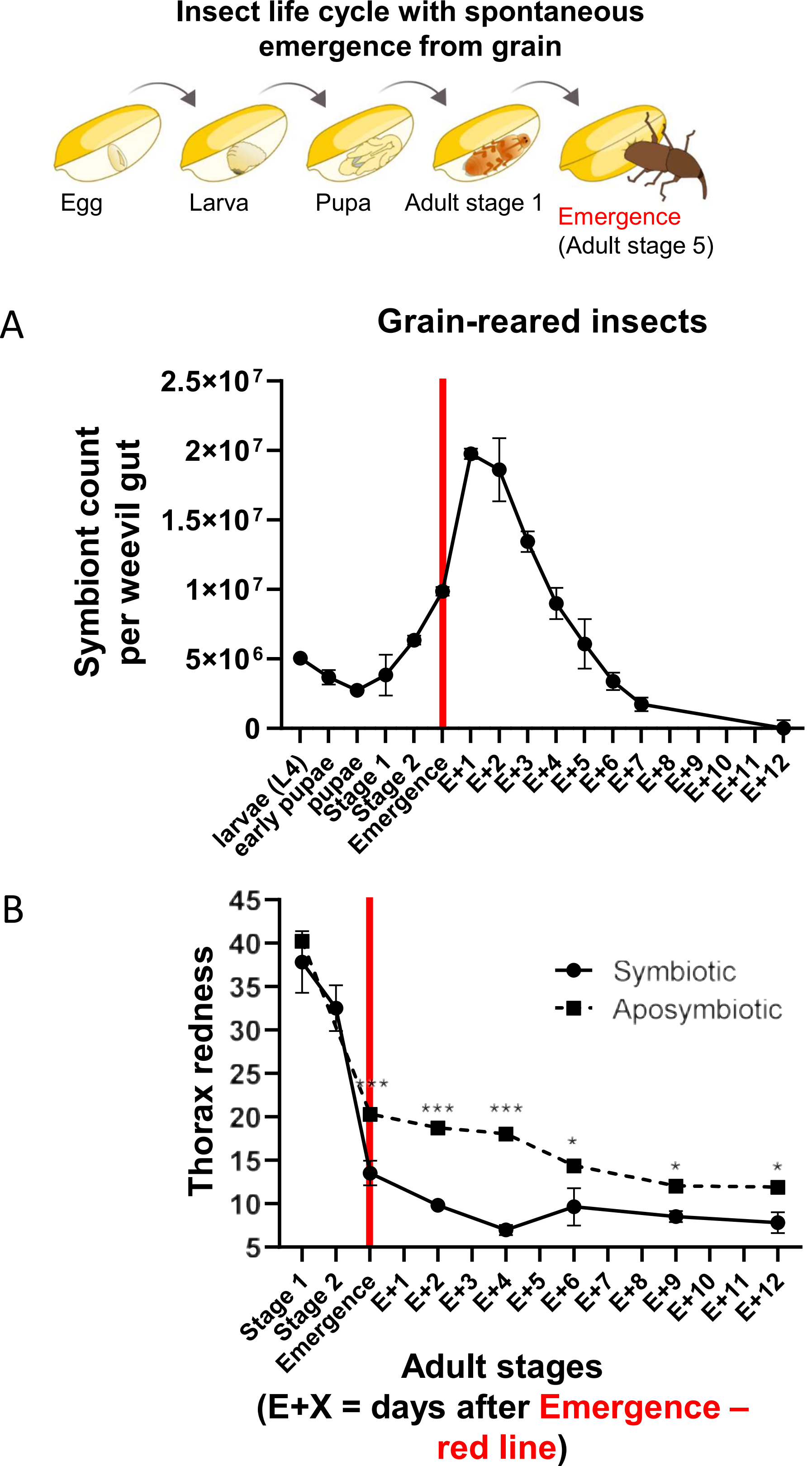

### Figure S2

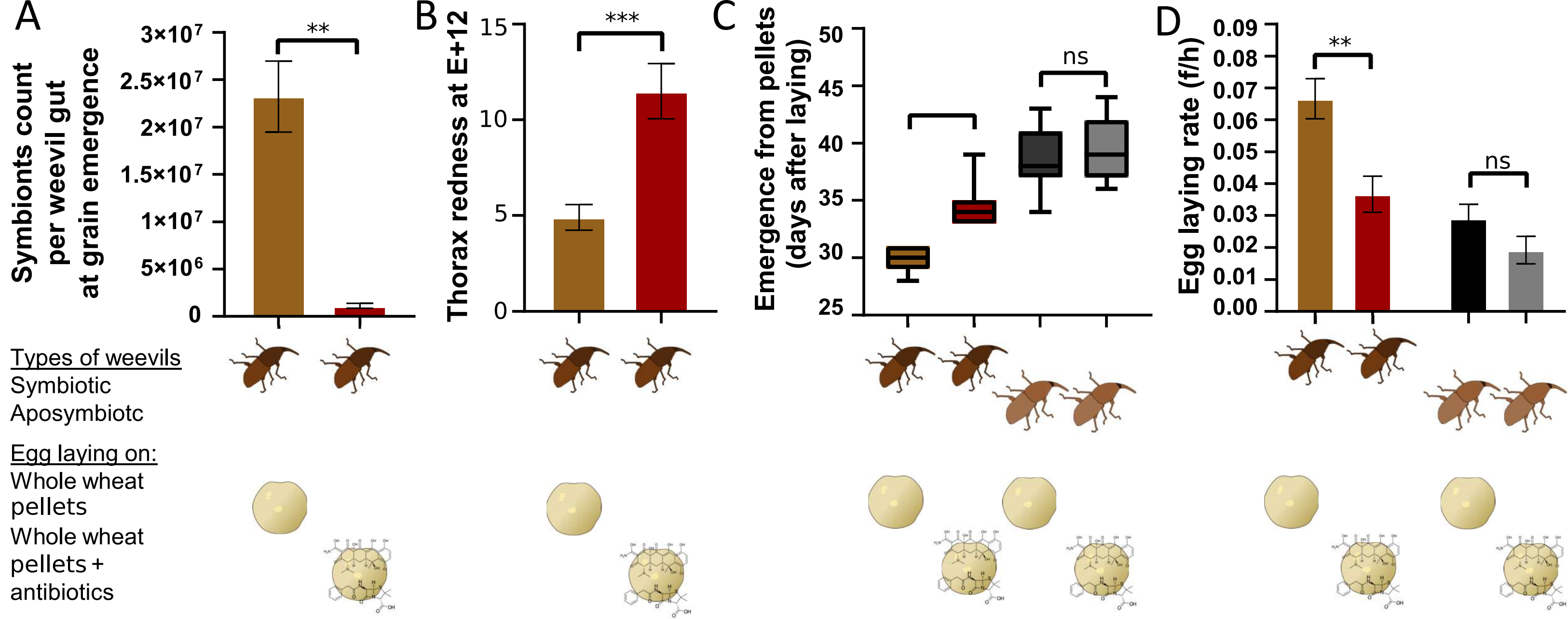

### Figure S3

A

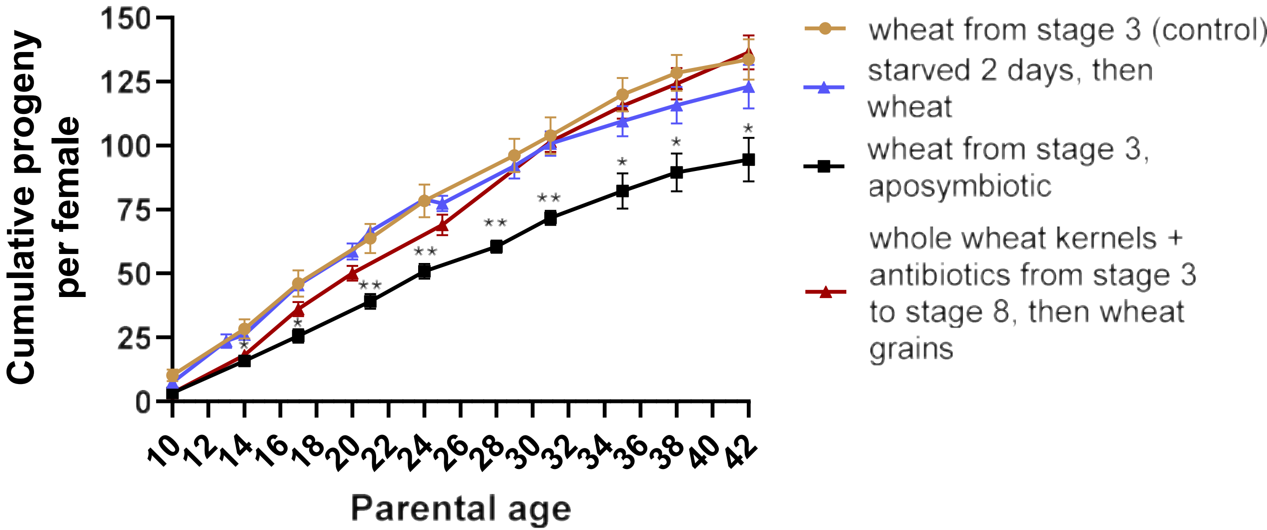


B

**Ovaries at stage 24**

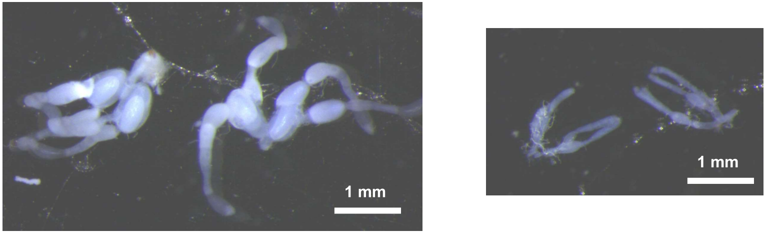


wheat from stage 3 (control)

starch diet

### Figure S4

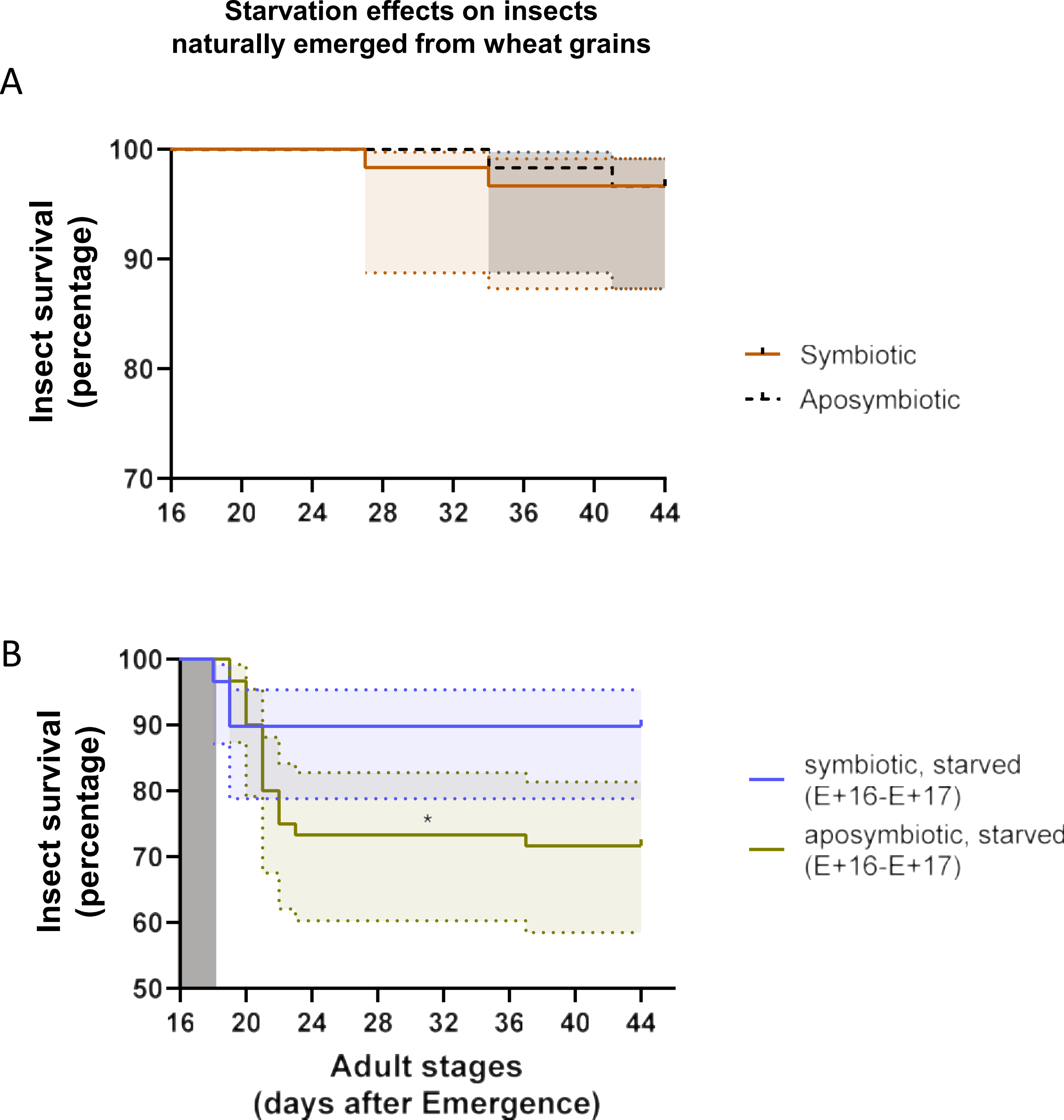

### Figure S5

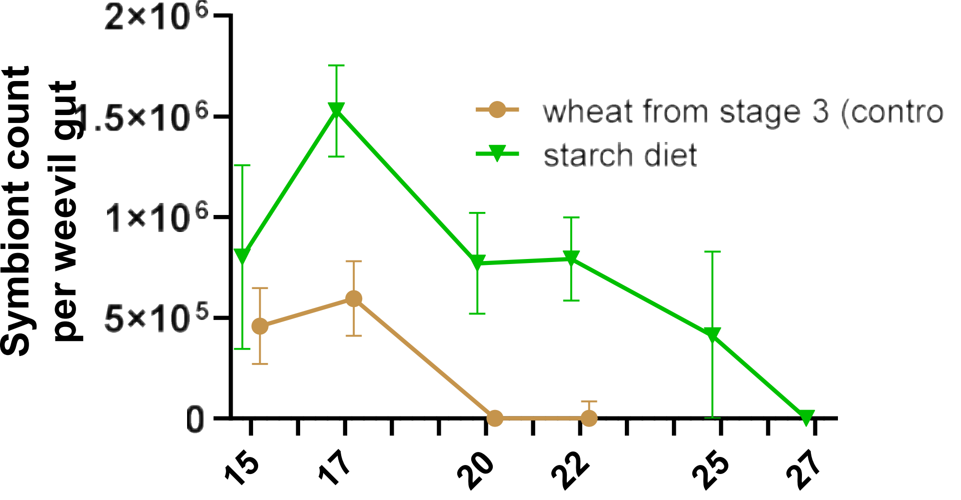

### Figure S6

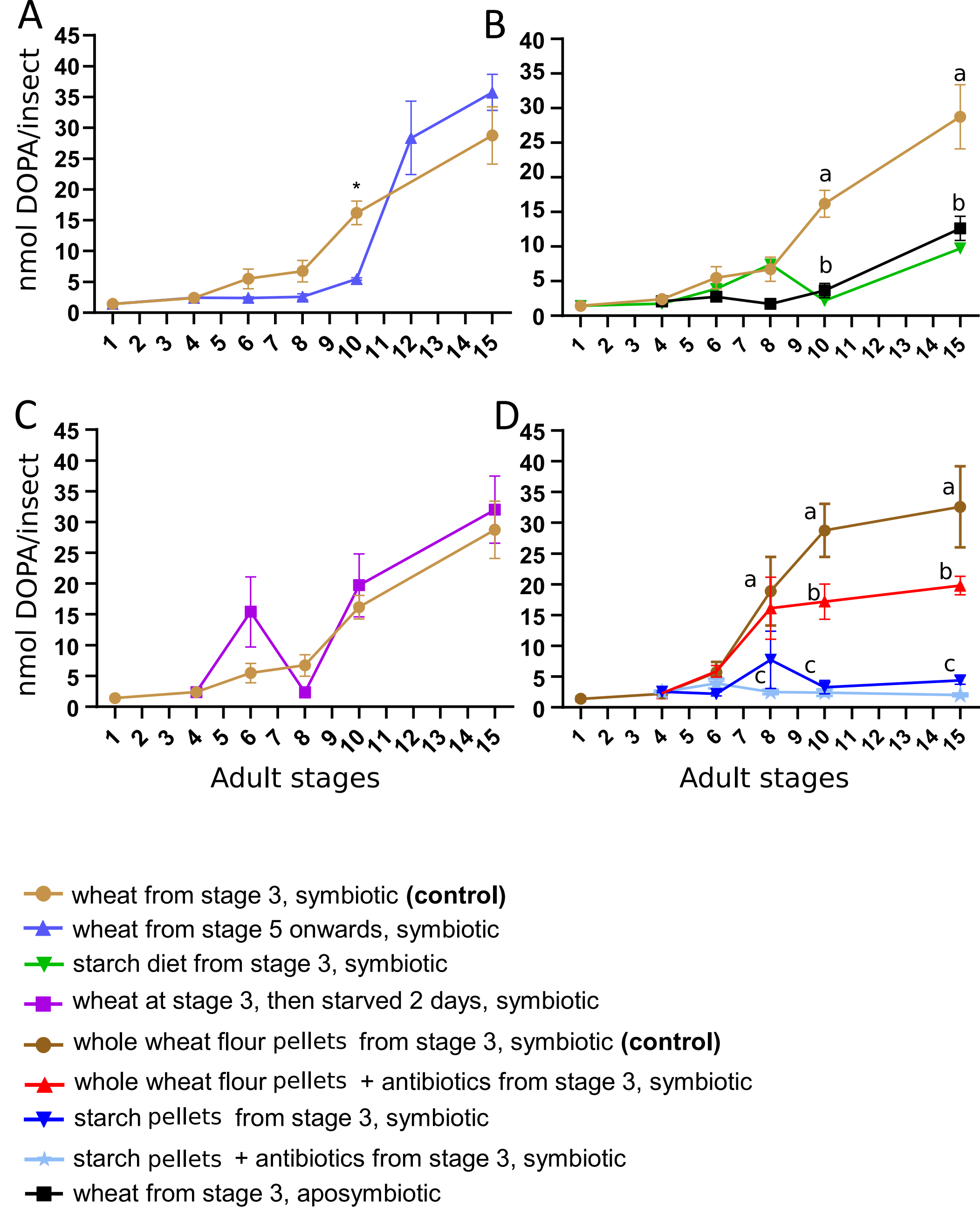
